# Gfastats: conversion, evaluation and manipulation of genome sequences using assembly graphs

**DOI:** 10.1101/2022.03.24.485682

**Authors:** Giulio Formenti, Linelle Abueg, Angelo Brajuka, Nadolina Brajuka, Cristo Gallardo, Alice Giani, Olivier Fedrigo, Erich D. Jarvis

## Abstract

**Motivation:** With the current pace at which reference genomes are being produced, the availability of tools that can reliably and efficiently generate genome assembly summary statistics has become critical. Additionally, with the emergence of new algorithms and data types, tools that can improve the quality of existing assemblies through automated and manual curation are required.

**Results:** We sought to address both these needs by developing gfastats, as part of the Vertebrate Genomes Project (VGP) effort to generate high-quality reference genomes at scale. Gfastats is a standalone tool to compute assembly summary statistics and manipulate assembly sequences in fasta, fastq, or gfa [.gz] format. Gfastats stores assembly sequences internally in a gfa-like format. This feature allows gfastats to seamlessly convert fast* to and from gfa [.gz] files. Gfastats can also build an assembly graph that can in turn be used to manipulate the underlying sequences following instructions provided by the user, while simultaneously generating key metrics for the new sequences.

**Availability and implementation:** Gfastats is implemented in C++. Precompiled releases (Linux, MacOS, Windows) and commented source code for gfastats are available under MIT license at https://github.com/vgl-hub/gfastats. Examples of how to run gfastats are provided in the Github. Gfastats is also available in Bioconda, in Galaxy (https://assembly.usegalaxy.eu) and as a MultiQC module [1] (https://github.com/ewels/MultiQC). An automated test workflow is available to ensure consistency of software updates.

**Supplementary information:** Supplementary data are available at Bioinformatics online.

## Introduction

In recent years we have witnessed an unprecedented increase in the number of publicly available genomes [2,3]. Thanks to advancements in genome sequencing and assembly [4], these genomes tend to be highly accurate and contiguous. Reference genomes are made available in publicly maintained archives such as Genbank by the US National Center for Biotechnology Information (NCBI, www.ncbi.nlm.nih.gov/genbank), the European Nucleotide Archive (ENA, www.ebi.ac.uk/ena/browser/home) by the European Bioinformatics Institute, the DNA Data Bank of Japan (DDBJ, www.ddbj.nig.ac.jp) or the China National Genebank (CNGB, https://db.cngb.org), or project-related repositories such as the Vertebrate Genomes Project (VGP) Genome Ark (https://vgp.github.io/) [4]. Assemblies are usually stored as collections of sequences representing either contigs (*i*.*e*. contiguous stretches of nucleotide sequences) or scaffolds (*i*.*e*. contigs separated by gaps of unknown sequence). The size of gaps can be approximately estimated (sized gaps) or unknown.

Sequence collections are generally stored in the popular FASTA format developed in 1985 [5]. In FASTA, each sequence is introduced by a “>” character followed by a header and a comment, and the sequence on newlines. Similar to FASTA, the FASTQ format was developed over two decades ago at the Wellcome Trust Sanger Institute [6] and later popularized by Illumina to store short read sequencing data with per-base quality information. More recently, the representation of biological sequences has been expressed under the conceptual framework of graph theory [7]. In a graph, genome assemblies can be represented as collections of sequences (nodes) linked by experimental evidence (edges). GFA is a popular format to store sequence data as graphs. GFA1, (http://gfa-spec.github.io/GFA-spec/GFA1.html) which was introduced in 2014 (http://lh3.github.io/2014/07/19/a-proposal-of-the-grapical-fragment-assembly-format), can be used to conveniently store and visualize [8] key features of sequence graphs, such as the product of an assembly [9], the representation of variation in genomes or overlaps between reads. Since the graph is not yet collapsed to a linear representation, many additional characteristics can be deduced. The GFA1 format consists of lines with tab-delimited fields. The first field defines the line type, which in turn defines additional required fields, followed by optional fields. Examples of line types are segments (S, usually a contig) and edges (L, usually an overlap between two contigs). GFA was later generalized to GFA2, which allows specifying an assembly graph in either less detail (e.g. only the topology of the graph) or in more detail (e.g. the multi-alignment of reads underlying each sequence) (https://github.com/GFA-spec/GFA-spec/blob/master/GFA2.md). Importantly, GFA2 introduces more line types to include gaps (G), allowing scaffolds (i.e. contig separated by gaps) to be represented.

As more and more reference genomes become available, a single fast, versatile tool that can compute assembly summary statistics from a variety of file formats is warranted. In the framework of the VGP, which aims to generate high-quality genome assemblies for all vertebrate species, we have developed and present here gfastats (short for graph-based *fa* statistics). By internally representing any input sequence (FASTA, FASTQ, GFA1/2) in a more general GFA2-like format, gfastats can efficiently compute accurate summary statistics. It further allows simultaneous manipulation of the assembly sequences, thereby potentially facilitating both automated assembly and manual curation.

## Results

For optimization purposes, gfastats is coded solely in C/C++, taking full advantage of object-oriented programming. In gfastats v1.2.0 (the version presented hereinafter), contigs (segments), edges and gaps are represented with classes, and so are the collections of paths through contigs and gaps that, taken together, represent a genome assembly (**Figure 1a**). Features of interest are represented using bed coordinates. Input includes any *fa* (fasta, fastq, gfa [.gz]) file. Since gfastats reads and stores any input in a gfa-like format, it allows the seamless conversion between different formats (fasta<>fastq<>gfa[.gz]). Inputs are processed on the fly to generate summary statistics.

**Figure 1.**
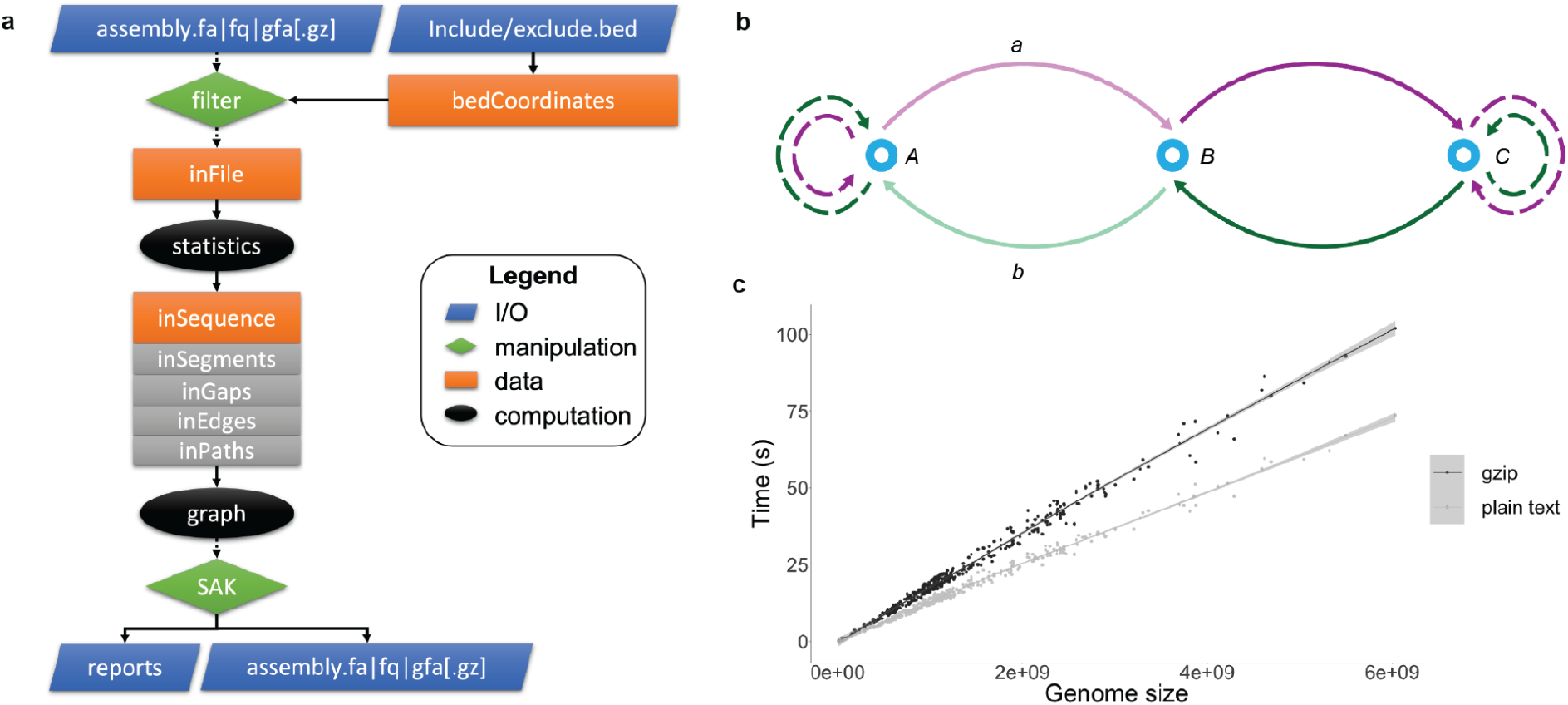
a) Schematic of gfastats workflow. Inputs (blue) include genome assemblies in fasta, fastq, gfa [.gz] formats and include/exclude lists as bed coordinate files for filtering (green). These are represented internally by multiple C++ classes (orange) and decomposed in their constituent elements (gray). Statistics can be generated for the assembly (black), or the assembly can be converted to a graph (black), to ease manipulation by the internal Swiss Army Knife (SAK; green). A variety of output can be generated, including summary statistics and new sequences in *fa* format. b) Internal bidirected graph representation of the input sequences. Segments/contigs (nodes, light blue) are connected by forward (purple) or backward (green) gaps (edges). Terminal nodes can optionally contain sequence gaps (dashed lines). So an assembly scaffold is a path in the graph (e.g. A → a → B → c → C). Sequence manipulation can be achieved using the internal SAK. For instance, the gap edges *a,b* connecting segment nodes *A,B* can be removed leading to a disconnected component with a starting gap *A*, and connected graph *A*-*B* with a terminal gap. Overlap edges can be similarly treated. c) Evaluation of gfastats runtime. Performance time is a function of genome size, with gfastats runtime increasing linearly. There is a small increase in time when handling gzip compressed files.

Gfastats computes a growing number of assembly/sequence metrics (**Figure 1a;** see **Supplementary Table 1** for a complete list and a comparison with other tools that provide assembly summary statistics). Metrics for each contig can also be generated. AGP (A Golden Path), BED coordinates and sizes of scaffolds, contigs and gaps can be conveniently outputted. Input can be filtered in a pre-processing step to include/exclude sequences or portions of them using scaffold lists or bed coordinate files. Sequences can be sorted, either according to a list or to other characteristics (name, length, etc.). Gfastats also allows homopolymer (de)compression, a feature increasingly useful when dealing with long reads.

Importantly, gfastats can convert AGP to GFA paths, allowing the output of any automated scaffolding tool such as Bionano Solve (https://bionanogenomics.com/downloads/bionano-solve/) SALSA2 [10] to be integrated in the graph. This avoids flatting the assembly graph to a linear sequence, making the integration of information from different sources and algorithms possible. In addition, since the assembly process is still imperfect, manual manipulation of contig and scaffold sequences is also needed. Genome assemblies often undergo a long process of curation, in which experts manually validate and correct the assembly using evidence from the raw data [11]. The process is not just laborious, but it also relies on file format specifications not adapted and not specifically designed for this purpose. By representing any input sequence as a graph, gfastats allows their manual manipulation. For instance, gfastats can build a bidirected graph representation of the assembly using adjacency lists, where each node is a segment, and each edge is a gap (**Figure 1b**). Canonical algorithms (e.g. Depth First Search) are used to walk the graph. In this case, the manipulation is achieved by the internal ‘swiss army knife’ (SAK) for genome assembly. SAK evaluates a set of basic sequential instructions, *i*.*e*. actions to be performed one-by-one on the graph to manipulate the sequence (*e*.*g*. join or split contigs, reverse complement sequence, etc.). Here, the representation of the assembly as a graph allows several operations to be performed (*e*.*g*. the removal of all trailing Ns from scaffolds by dropping all terminal gap edges). Once all instructions for the SAK are processed, metrics are updated and returned, allowing evaluation of the revised assembly. The filtered and/or manipulated input can also be outputted in any *fa* format, thereby generating new sequences.

Testing on a 2.8 GHz Quad-Core Intel Core i7 using 370 genome assemblies from the VGP shows that gfastats can compute all summary statistics in less than a minute for genome assemblies of size up to 4 Gbp in O(N) time (**Figure 1c**). Assembly manipulation comes with minimal overhead.

## Discussion and future perspectives

As graph representations of genome assemblies become more popular, effective tools that make assembly graph storage, analysis and manipulation easily accessible become necessary. While a few libraries already exist to deal with GFA files [12] (https://github.com/lh3/gfatools), they do not make FASTA and GFA fully interoperable, and do not directly allow their seamless manipulation. The design of gfastats addresses this need in a modular framework, allowing new features to be readily implemented. Potential new features include: file indexing to test multiple hypotheses with minimal runtime overhead, pattern search, sequence soft/hard-masking, and new instructions to the SAK. Additional FASTA and GFA statistics can also be introduced based on the needs of the genomics community. Importantly, gfastats introduces a whole new conceptual framework for assembly manipulation where the results of automated algorithms or manual curation be integrated in a single file format and can be expressed in a human-readable set of instructions for the SAK, which also conveniently acts as a log of the changes that were applied during the process.

## Author contributions

G.F. implemented gfastats with contributions from A. G., A. B., L. A., N. B. and C. G. A. G., L. A, N. B. performed the comparative analyses. N. B. evaluated runtime performances using the VGP genomes. A. B. implemented the automated test workflow and homopolymer compression. C. G. implemented gfastats in Conda and Galaxy. A. G. contributed to the conceptual development. G. F. conceived the study and wrote the manuscript, with contributions from O. F. and E. D. J. All authors reviewed and approved the manuscript.

## Acknowledgments

We thank Björn Grüning for helping with the implementation of gfastats in Conda and Galaxy.

---

**Supplementary table 1:** Typical *fa* operations computed by gfastats for scaffolds, contigs (nodes), gaps (edges). Several metrics are missing from popular summary statistics tools, e.g. QUAST [13], SeqKit [14] and Bandage [8]. AuN, area under the curve.

